# Arginine citrullination at the C-terminal domain controls RNA polymerase II transcription

**DOI:** 10.1101/216143

**Authors:** Priyanka Sharma, Antonios Lioutas, Narcis Fernandez-Fuentes, Javier Quilez, José Carbonell-Caballero, Roni H.G. Wright, Chiara Di Vona, François Le Dily, Roland Schüller, Dirk Eick, Baldomero Oliva, Miguel Beato

## Abstract

**Highlights:**

- Peptidyl arginine deiminase 2 (PADI2) citrullinates arginine1810 (cit1810) present at carboxy-terminal domain of RNA polymerase II (RNAP2-CTD).
- PADI2 and R1810 of RNAP2-CTD regulate transcription and proliferation of breast cancer cells.
- Absence of cit1810 at RNAP2-CTD leads to RNAP2 accumulation at proximal promoter regions.
- Cit1810 at RNAP2-CTD facilitate interaction with P-TFEb complex.

**SUMMARY:** The post-translational modification of key residues at the carboxy-terminal domain of RNA polymerase II (RNAP2-CTD), coordinates transcription, splicing, and RNA processing by modulating its capacity to act as a landing platform for a variety of protein complexes. Here, we identify a new modification at the CTD, the deimination of arginine and its conversion to citrulline by peptidyl arginine deiminase 2 (PADI2), an enzyme that has been associated with several diseases including cancer. We show that among PADI family members, only PADI2 citrullinates R1810 (Cit1810) at repeat 31 of the CTD. Depletion of PADI2 or loss of R1810 result in accumulation of RNAP2 at transcription start sites, reduced gene expression and inhibition of cell proliferation. Cit1810 is needed for interaction with the P-TEFb (positive transcription elongation factor b) kinase complex and for its recruitment to chromatin. In this way, CTD-Cit1810 favors RNAP2 pause release and efficient transcription in breast cancer cells.

## INTRODUCTION

In mammals, the RNA polymerase II carboxy-terminal domain (RNAP2-CTD) comprises 52 heptapeptide repeats, the first half of which (1-27) exhibit the consensus repeat sequence Y_1_S_2_P_3_T_4_S_5_P_6_S_7_, whereas the second half (28-52) contains deviations from this consensus (Buratowski et al., 2009; Corden et al., 2013). Post-translational modification of the key residues at RNAP2-CTD dictate recruitment of protein complexes, that influence transcription elongation and the processing of the nascent transcripts (Jeronimo et al., 2016; Saldi et al., 2016; Harlen et al., 2017). The CTD is evolutionary conserved and dynamic phosphorylation of Y_1_, S_2_, T_4_, S_5_, and S_7_ mediates selective recruitment of protein complexes that modulate various phases of transcription (Eick et al., 2013; Zaborowska et al., 2016; Shah et al., 2018). Systematic studies using genetics and proteomics showed that phosphorylation of S5 and S2 is the most frequent modification and contributes to transcription efficiency (Schüller et al., 2016; Corden, 2016). However, modifications in non-consensus repeats have expanded the functional role of the CTD code (Voss et al., 2015; Dias et al., 2015) and recent work has focused on methylation of arginine 1810 (R1810) at repeat 31. Its asymmetrical dimethylation (me2a) by the methyltransferase CARM1 (or PRMT4) inhibits the expression of small nuclear RNAs (snRNAs) and nucleolar RNA (snoRNA) genes in human cells (Sims et al., 2011). This reaction is inhibited by phosphorylation of CTD serine residues, suggesting that it occurs before transcription initiation. In contrast, symmetric dimethylation (me2s) of R1810 by PRMT5 leads to recruitment of the survival of motor neuron protein (SMN) and to the interaction with senataxin, that enhances transcriptional termination (Zhao et al., 2016). The functional significance of dynamic post-translational deimination of arginine residues in pathophysiological conditions (Slade et al., 2014; Tanikawa et al., 2018) prompted us to investigate whether this modification occurs at R1810 of RNAP2-CTD and its possible implication in transcription regulation.

Citrullination is a deimination of protein-embedded arginine, which is converted to the non-coded amino acid citrulline (Van et al., 2000; Fuhrmann et al., 2015). Citrullination leads to a reduction in hydrogen-bond formation, affects histone–DNA interactions and influences the chromatin organization. Citrullination also increases the hydrophobicity of proteins that affect the protein folding ability and therefore the functional activity of proteins (Vossenaar et al., 2003; Tanikawa et al., 2018). This reaction is catalyzed by enzymes called peptidyl arginine deiminases (PADIs), which have been associated with diverse disease conditions such as thrombosis, prion disease, neurological disorders, autoimmune disease and cancer (Witalison et al., 2015; Gyorgy et al., 2006; Vossenaar et al., 2003; Baka et al., 2012). Among PADI family members, PADI2 is the most widely expressed isoform and is also overexpressed in breast cancer, where it regulates mammary carcinoma cell migration (Mohanan et al., 2012; Cherrington et al., 2012; Horibata et al., 2017). Citrullination of core histones has been related to the gene expression, DNA damage responses and pluripotency (Sharma et al., 2012; Tanikawa et al., 2012; Christophorou et al., 2014), although the underlying mechanisms are largely unknown.

RNAP2-mediated gene expression starts with binding to the gene promoters of basal transcription factors that recruit RNAP2 to form the transcription pre-initiation complex. Shortly after transcription initiation, RNAP2 pauses 30-50bp downstream of the transcription start sites (TSS), and requires the activation of P-TEFb (positive transcription elongation factor b) kinase complexes to continue with the productive elongation (Marshall et al., 1995; Adelman et al., 2012; Jonkers et al., 2015). Promoter-proximal pausing affects the expression of many genes but is more prominent for highly expressed genes in responses to developmental and environmental stimuli (Zeitlinger et al., 2007; Core et al., 2008; Gilchrist et al., 2010; Day et al., 2016). Recently, RNAP2 pausing was found to inhibits transcriptional initiation, indicating that paused RNAP2 first needs to be released in order to allow a new cycle of transcription inititation (Shao et al., 2017; Gressel et al., 2017). However, despite the strong evidence of RNAP2 pausing, the nature of paused RNAP2 to allow efficient transcription still remains unclear.

Here, we report the discovery that PADI2 citrullinates the R1810 (cit1810) at RNAP2-CTD. The absence of PADI2 mediated cit1810 widely affects transcription and cell proliferation in breast cancer cells. PADI2 occupancy increases with the level of gene transcription. Further, we found that replacing wild-type RNAP2 with the R1810A mutant compromises transcription, reduces interaction with the P-TEFb complex and leads to accumulation of RNAP2 on the proximal promoter of PADI2 dependent genes. Thus, citrullination of R1810 facilitates interaction with the P-TEFb complex favoring RNAP2 pause release and promoting transcription of cell cycle genes and cell proliferation of breast cancer cells.

## RESULTS

### Citrullination of R1810 at RNAP2-CTD

Two arginine residues within non-consensus repeats in human RNAP2-CTD, R1603 and R1810, are conserved in vertebrates. Recently, R1810 within repeat 31 was found to be asymmetrically (Sims et al., 2011) or symmetrically (Zhao et al., 2016) dimethylated, leading to either reduced expression of snRNAs and snoRNA or to efficient transcription termination, respectively. To examine the possibility that R1810 at RNAP2-CTD could be citrullinated in cells, we immunoprecipitated nuclear extracts from the luminal breast cancer cell line T47D (Truss et al., 1995) with a citrulline antibody followed by western blot with an antibody to RNAP2. We found two specific bands migrating as the non-phosphorylated (IIA) and phosphorylated (IIO) forms of the large subunit POLR2A of RNAP2. The IIO band reacted preferentially with α-citrulline compared to the IIA band of RNAP2 (**Figure S1A**). We raised a polyclonal antibody against a 13 residues peptide centered on R1810, which was replaced by citrulline (**Figure 1A top**). This antibody (α-Cit1810) was specific, as it reacted with the citrullinated peptide, but not with the wild-type, methylated (me2aR1810) or phosphorylated (S2 or S5) peptides (**Figure S1B-C**) and mainly recognized the phosphorylated form of RNAP2 in western blots of nuclear extracts from T47D cells (**Figure 1A**). RNAP2–Cit1810 was detected in several other cancer cell lines derived from the breast (BT474, SKBR3, MCF7), brain (SK-N-SH, T98G), and cervix (HeLa), but not in untransformed breast epithelial cells (MCF10A) or in normal fibroblasts (**Figure S1D**).

**Figure 1 (With related Figure S1).**
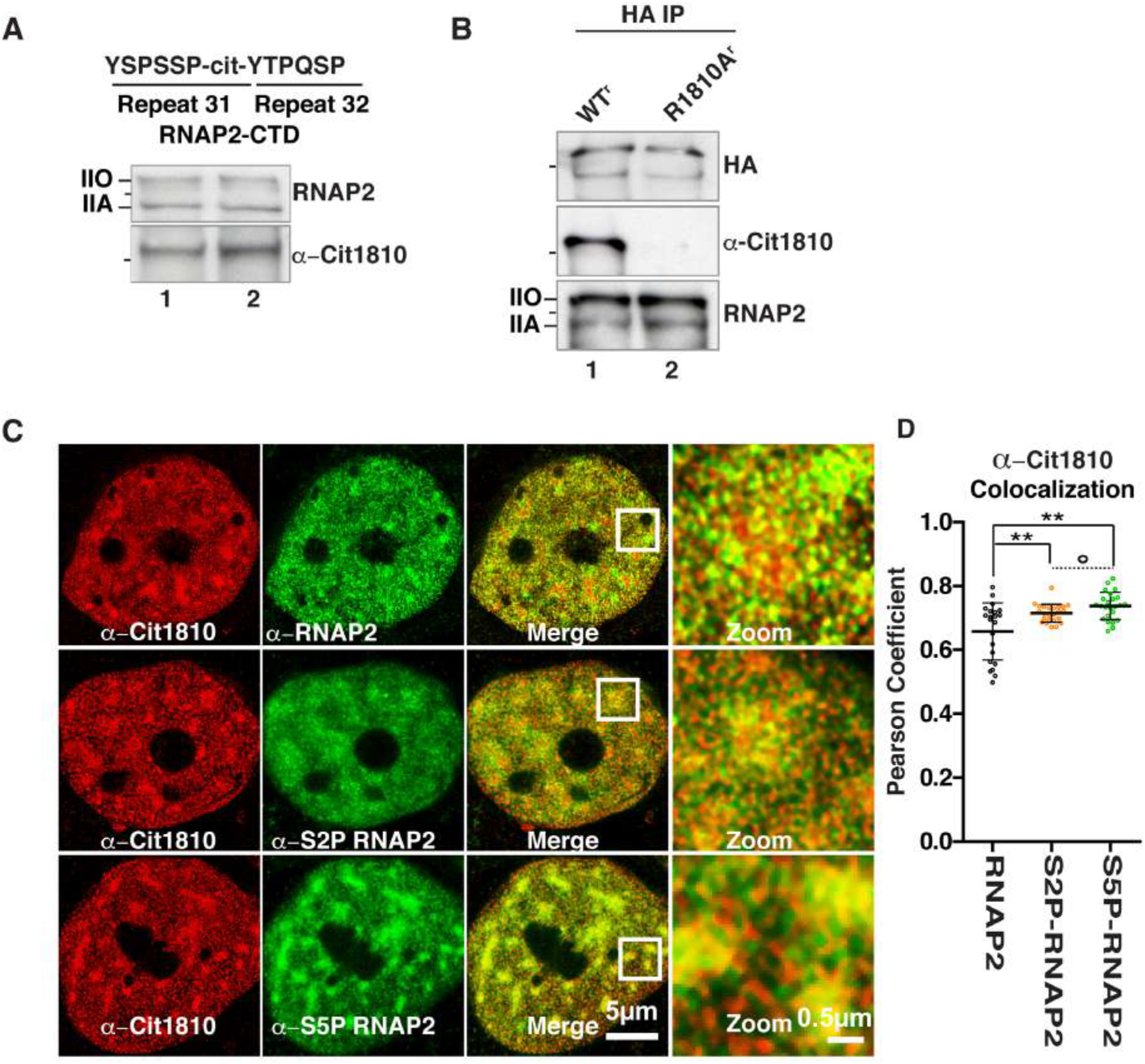
Citrullination of R1810 at RNAP2-CTD. (**A**). *Top:* the epitope within repeats 31/32 of the CTD domain of RNAP2 used to generate α-Cit1810. *Bottom:* duplicated western blot of T47D nuclear extract with α-Cit1810 and α-total-RNAP2. Line on the left mark the migration of the 250 kDa size marker. (**B**). Extracts from T47D cells expressing α-amanitin resistant WT^r^ or R1810A^r^ mutant of RNAP2 were precipitated with α-HA antibody and probed with α-Cit1810 or α-RNAP2. (**C**). Representative super-resolution images of T47D cells immunostained with α-Cit1810 (red) in combination with α-total-RNAP2 (green), α-S2P-RNAP2 (green) and α-S5P-RNAP2 (green). (**D**) Plot representing the mean Pearson correlation coefficient of individual cells for α-Cit1810-RNAP2 with α-total-RNAP2 (n=22), α-S2P-RNAP2 (n=24) and α-S5P-RNAP2 (n=24); values presented as the mean ± SEM. ** p-value < 0.005; ° p-value > 0.05.

To validate that R1810 is citrullinated, we transiently transfected T47D cells with an α-amanitin resistant HA-tagged wild-type (WT^r^) RNAP2 or with a R1810A^r^ mutant of RNAP2, followed by α-amanitin treatment to deplete the endogenous RNAP2 (**Figure S1 E-F**). Precipitation with anti-HA antibody followed by western blot showed that the WT^r^ RNAP2, but not the R1810A^r^ mutant, reacts with α-Cit1810 (**Figure 1B**). In superresolution immunofluorescence images of T47D cells, α-Cit1810 decorated bright clusters overlapping with RNAP2, preferentially in its S2 or S5 phosphorylated forms (**Figure 1C-D**). Thus, R1810 is citrullinated in cells prevalently on the phosphorylated actively transcribing form of RNAP2.

### Citrullination of R1810 by PADI2

In a search for the responsible enzyme, we found that T47D cells express only *PADI2* and *PADI3* (**Figure 2A**) among family members. Depletion of PADI2 but not PADI3 reduces R1810 citrullination (**Figure 2B, Figure S2A**). In MCF7 breast cancer cells that express *PADI2* and *PADI4* (Cuthbert et al., 2004; Sharma et al., 2012), depletion of *PADI2* but not PADI4 reduced R1810 citrullination (**Figure S2B-C**). To test whether PADI2 acts directly on the RNAP2-CTD we incubated recombinant PADI2 with either a recombinant GST-N-CTD (repeat 1-25.5, including R1603) or with GST-C-CTD (repeat 27-52, including R1810) (**Figure 2C**, *left panel*). PADI2 citrullinates R1810 in the C-CTD much more efficiently than R1603 in N-CTD (**Figure 2C**, *right panel*). Thus, PADI2 is the enzyme responsible for citrullination of R1810 in breast cancer cells

**Figure 2 (With related Figure S2-4).**
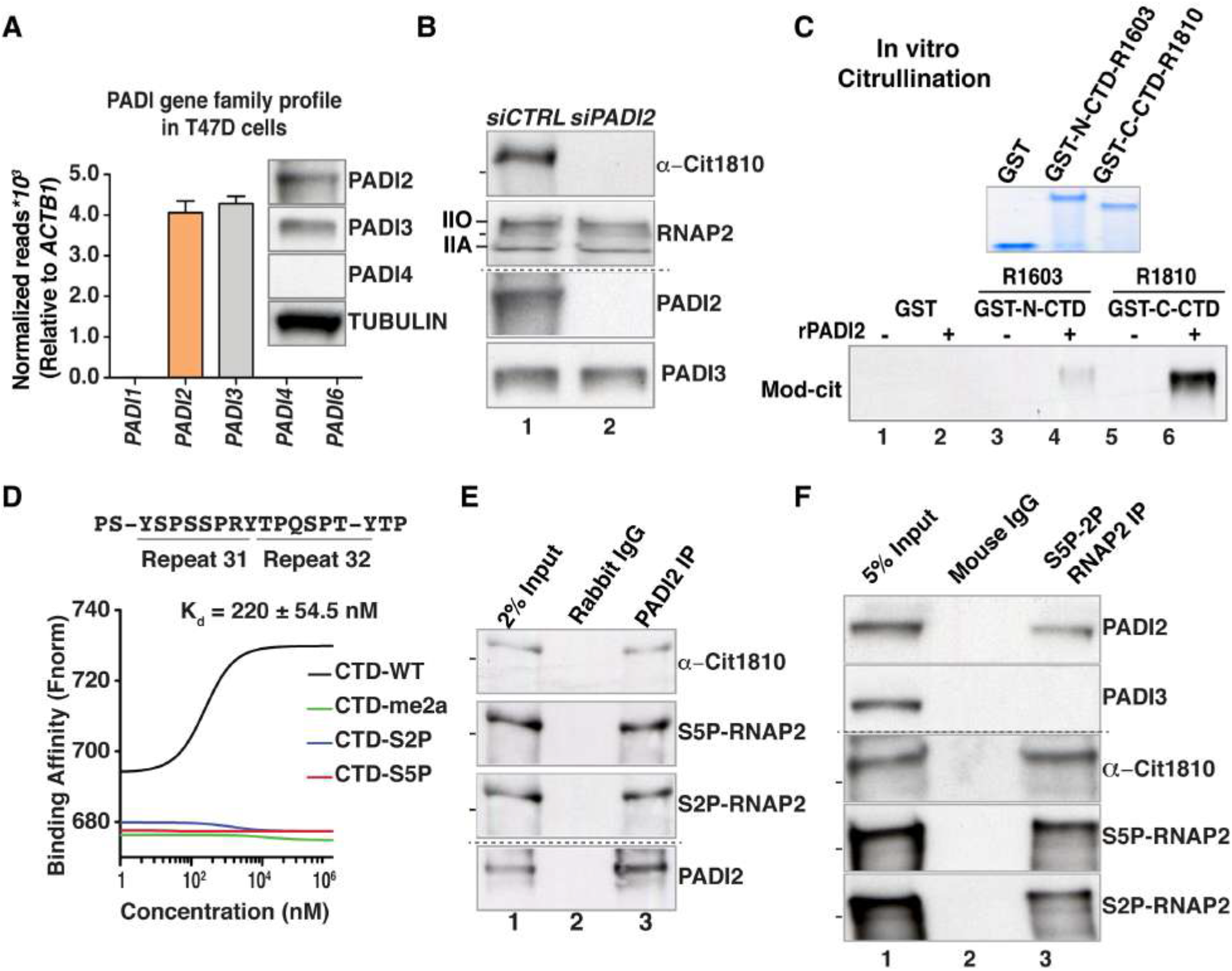
PADI2 citrullinates R1810 at RNAP2-CTD. (**A**) Bar plot showing the normalized reads of *PADI* gene family members from two RNA-sequencing experiments performed in T47D cells. The normalized reads are represented relative to *Actb1* gene. Values are means ± SEM. Inset-western blots performed on T47D nuclear extract with antibodies to PADI2, 3, 4 and Tubulin. (**B**) Nuclear extracts from T47D cells transfected with siRNA control (si*CTRL*) or siRNA against PADI2 (si*PADI2*) are probed with α-Cit1810, α-total-RNAP2, α-PADI2 and α-PADI3. (**C**) *Top:* Coomassie blue staining of SDS-PAGE with recombinant GST-tagged, GST-N-CTD and GST-C-CTD proteins used for the citrullination assay. *Bottom: In vitro* citrullination immunoblot with or without recombinant PADI2 (rPADI2) using as substrate the N-terminal half of the CTD containing R1603 (*lanes 3 & 4*) or the C-terminal half containing R1810 (*lanes 5 & 6*), both linked to GST. As a control, GST was also tested (*lanes 1 & 2*). (**D**) Microscale thermophoresis assay showing the affinity of recombinant PADI2 for the indicated wild -type and modified CTD-RNAP2 peptides encompassing the R1810. Y-axis represents the binding affinity as normalized fluorescence (Fnorm, see methods). (**E-F**) Immunoprecipitation with (**E**) α-PADI2 (F) α-S2P/S5P RNAP2 or non-immune mouse or rabbit IgG of T47D extracts followed by western blot with the indicated antibodies.

The affinity of PADI2 for the unmodified R1810 peptide measured by microscale thermophoresis (Jerabek et al., 2011) is K_d_=220±54.5 nM, whereas peptides phosphorylated at S2 or S5 were not bound (**Figure 2D**), suggesting that the observed S2/S5 phosphorylation in R1810 citrullinated CTD must occur outside of repeats 31 and 32. In co-immunoprecipitation experiments using T47D and MCF7 cells extracts, PADI2 but neither PADI3 or nor PADI4 interacted with RNAP2 (**Figure S2D-E**). Similarly, an antibody against PADI2 precipitated Cit1810-RNAP2, along with S5P- and S2P-RNAP2 (**Figure 2E, Figure S2F**). Monoclonal antibodies against RNAP2 phosphorylated at S2 and S5 (see methods) precipitated Cit1810 RNAP2 as well as PADI2 but not PADI3 (**Figure 2F**). In T47D nuclear extracts fractionated using size exclusion chromatography PADI2 eluted along with phosphorylated RNAP2 in the high molecular weight fractions, whereas PADI3 eluted in lower molecular weight range (**Figure S2G**). Finally, triple labeling immunofluorescence microscopy showed that PADI2 co-localizes with Cit1810-RNAP2 and with S2P-RNAP2 (**Figure S2H**). These observations support the association of PADI2 with Cit1810-RNAP2 that is engaged in transcription. To explore whether PADI2 was also expressed in samples from cancer patients, we analyzed the expression of *PADI* gene family across cancer cohorts and found that only PADI2 is overexpressed in breast and other cancers (**Figure S3A-E**) and that *PADI2* overexpression correlates with poor prognosis (**Figure S4A-E**).

### Citrullination of R1810 regulates transcription and cell proliferation

We next performed mRNA sequencing in control and PADI2 depleted T47D cells (**Figure S5A**). Strikingly, in global differential expression (DEseq) analysis PADI2 knockdown affected the expression of over 4,000 genes; downregulated (2,186) and upregulated (2,141) (**Figure 3A, Figure S5B**). Gene ontology analysis of the down-regulated genes revealed enrichment in RNAP2-mediated transcription and cell proliferation (**Figure S5C**). Reduced expression was validated by RT-qPCR for several genes including *SERPINA6, c-MYC*, and *HMGN1* genes, while control genes *GSTT2* and *LRRC39* were not affected (**Figure 3B, Figure S5D**). Depletion of CARM1 or PRMT5, which catalyze dimethylation of R1810 (Sim et al., 2011; Zhao et al., 2016), did not affect the expression of PADI2-dependent genes (**Figure S5E-F**), indicating that R1810 dimethylation and citrullination affect the expression of different sets of transcripts.

**Figure 3 (With related Figure S5).**
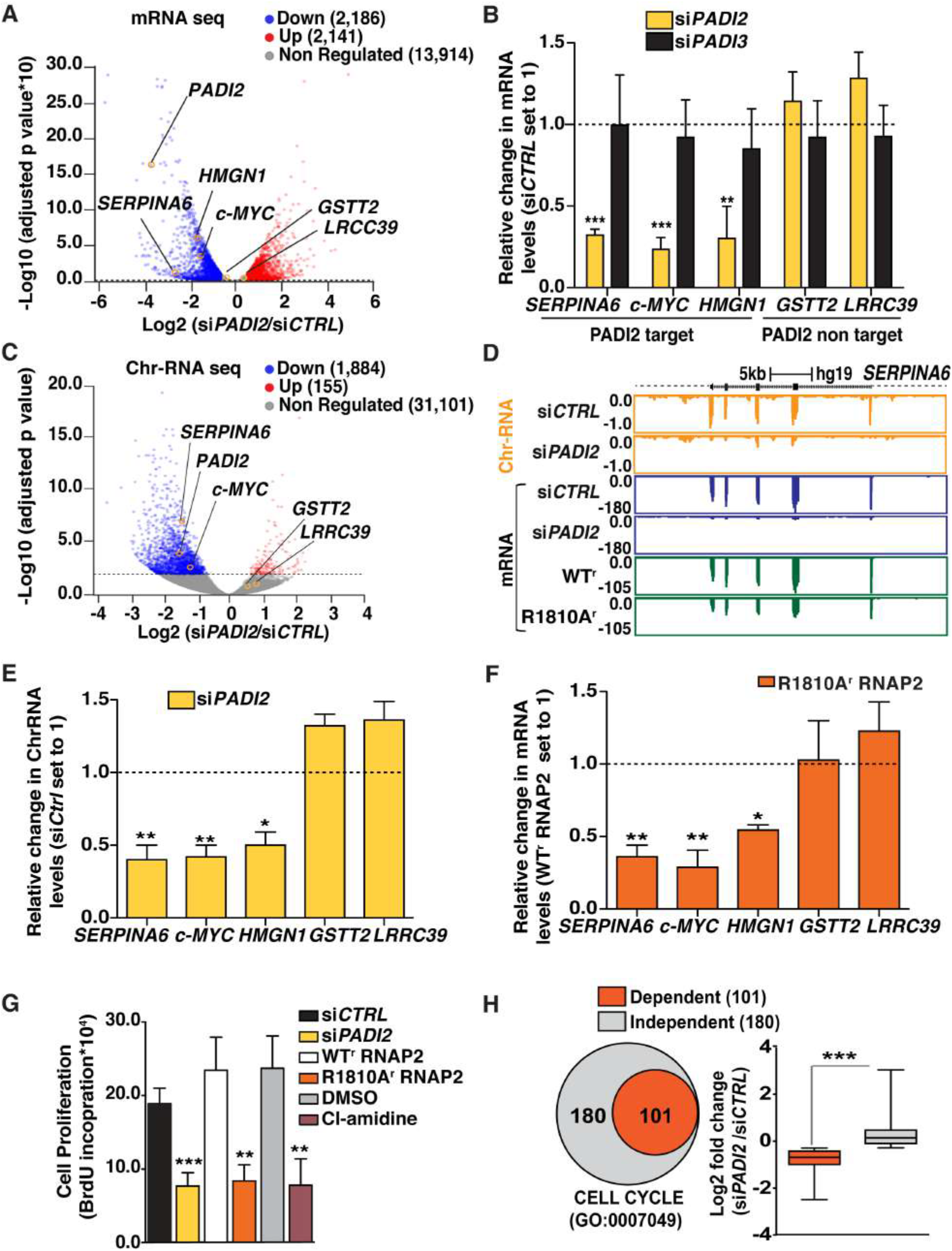
PADI2 mediated cit1810 at RNAP2-CTD regulates transcription and cell proliferation in breast cancer cells. (**A**) Volcano plot showing genome-wide mRNA changes after PADI2 depletion from biological replicates. The X-axis represents log2 expression fold changes (FC) and the Y-axis represents the adjusted p-values (as −log10). Differentially expressed genes (FC > 1.5 or <1/1.5 and p value < 0.01) are shown, the positions of *PADI2* and genes used for validation are also indicated. (**B**) Quantitative RT-qPCR validation in T47D cells transfected with si*CTRL*, si*PADI2* and si*PADI3*. Changes in mRNA levels were normalized to *GAPDH* mRNA. Data represent mean ± SEM of at least three biological experiments as in other plots in the figure. (**C**) Volcano plot showing genome-wide chromatin associated RNAs changes before and after PADI2 depletion from independent replicates. The X-axis represents the log2 fold changes (FC) and the Y-axis represents the adjusted p values (−log10). The dotted line indicates the cutoff p value < 0.01. Differential expressed genes (FC > 1.5 or 1/1.5 and the p value < 0.01) are shown. (**D**) Browser snapshots of *SERPINA6* gene in T47D cells showing chromatin associated RNAs sequencing profile (orange) and mRNA sequencing profiles after *PADI2* knockdown (blue) and expressing WT^r^ and R1810^r^ mutant form of RNAP2 (Green). Scale is indicated on the top of the gene. (**E**) Quantitative RT-qPCR on chromatin associated RNA (ChrRNA) in T47D cells transfected with si*CTRL*, si*PADI2* RNAs. Data normalized to *GAPDH* ChrRNA expression level. (**F**) T47D cells expressing only α-amanitin resistant HA tagged WT^r^ or R1810A^r^ mutant form of RNAP2 were used for quantitative RT-qPCR of mRNA from PADI2 dependent genes (*SERPINA6, c-MYC* and *HMGN1*), and for control genes (*GSTT2* and *LRRC39*). (**G**) Cell proliferation of T47D cells in the absence (si*CTRL* or DMSO) or presence of PADI2 depletion (si*PADI2*), PADI2 inhibitor (Cl-amidine in DMSO) and of cells expressing α-amanitin resistant HA tagged WT^r^ and R1810A^r^ mutant form of RNAP2. (**H**) ***Left*:** Venn diagram showing the set of genes related to cell cycle (GO:0007049 (n= 315, Out of them 281 genes expressed in T47D cells) downregulated in T47D cells after PADI2 depletion (PADI2 dependent) versus PADI2 independent gene. ***Right*:** Box plot showing log2 fold change (si*CTRL*/si*PADI2*) for PADI2 dependent and independent cell cycle genes. Each box in the panel represents the interquartile range; Whisker extends the box to the highest and lowest values, horizontal lines indicate the median value. Dependent genes showed significant lower mRNA levels than independent genes (***p-value < 0.0001, calculated by Wilcoxon-Mann-Whitney test).

To explore the direct effect of PADI2 on nascent transcription, we performed chromatin-associated RNA sequencing (ChrRNA-seq, Nojima et al., 2016) in control and PADI2 depleted cells. We found that ~2,000 transcripts were significantly affected by the PADI2 knockdown, and the majority of them were downregulated (1,884, **Figure 3C-D, Figure S5G**). ChrRNA-seq changes were validated on PADI2-dependent genes by RT-qPCR (**Figure 3E**). We conclude that PADI2 is required for efficient transcription and that upregulation of mRNAs upon PADI2 depletion may be a consequence of down-regulation of transcription relevant genes, although we cannot exclude citrullination of other PADI2 substrates. To support this conclusion, we performed mRNA sequencing in T47D cells expressing only the α-amanitin resistant HA-tagged WT^r^ or the R1810A^r^ mutant form of RNAP2. We found 1,392 down-regulated genes in cells expressing R1810A^r^ mutant RNAP2, of which 939 (67.4%) were also dependent on PADI2. We confirmed this finding by RT-qPCR of a PADI2-dependent *SERPINA6, c-MYC*, and *HMGN1* and control genes *GSTT2* and *LRRC39* (**Figure 3F**). Thus, PADI2 and R1810 are required for efficient transcription.

Since many PADI2-dependent genes are related to cell proliferation, we monitored T47D cell proliferation after PADI2 depletion (si*PADI2*), inhibition with Cl-amidine, or in cells expressing only the R1810A^r^ mutant of RNAP2. In all cases, we found a significant reduction of cell numbers (**Figure 3G**). PADI2-depleted cells were arrested at the G1 phase of the cell cycle (**Figure S5H**), as expected given the downregulation of genes critical for G1 phase progression including *CCND1, PLK1* (**Figure 3H, Figure S5I**).

### PADI2 is enriched on active genes

ChIP-seq of PADI2 in T47D cells showed that 60% of chromatin-bound PADI2 was localized over protein-coding gene sequences, within 3kb upstream of the TSS (transcription start site) and 3kb downstream of the TTS (transcription termination site) (q value ≤0.005) (**Figure 4A**, *left panel*)). The highest enrichment (2.5-fold) was found in the coding exons, followed by the 3kb region downstream of the TTS (1.6-fold) (**Figure 4A**, *Right panel*)). Overall, PADI2 occupancy overlapped with RNAP2 binding measured by ChIP-seq (**Figure 4B**). To explore whether PADI2 binding is related to transcription, we separated genes in 4 quartiles according to their transcription level: high (100-95%), medium (95%-50%), low (last 50%) and silent (non-significant expression) (Baranello et al., 2016). We found that RNAP2 and PADI2 occupancy increased in parallel with the gene expression levels (**Figure 4C, Figure S6A**), supporting a role of PADI2 in transcription.

**Figure 4 (With related Figure S6).**
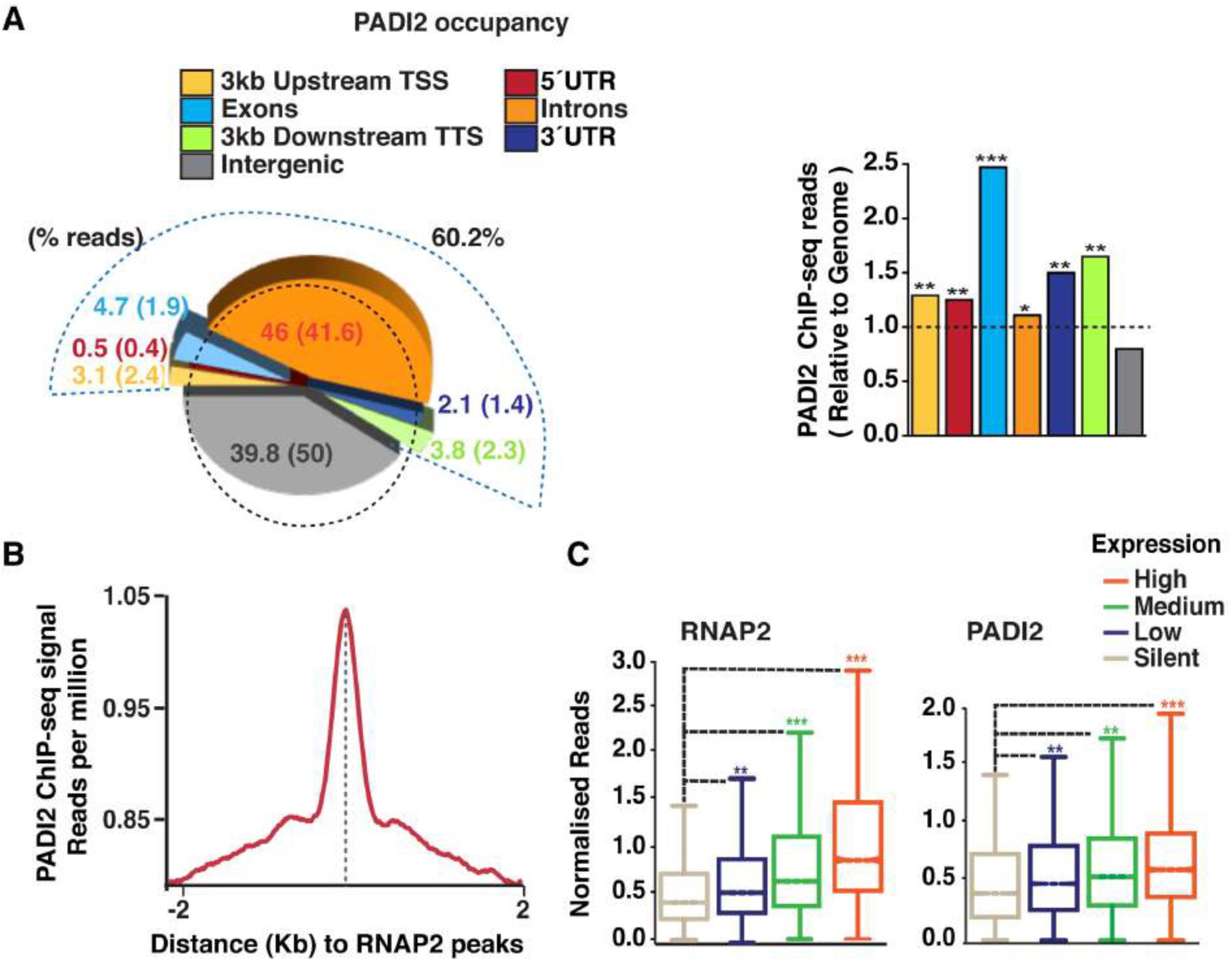
PADI2 occupancy on active genes. (**A**) *Left*: Spie chart showing the distribution of PADI2 ChIP-seq peaks over various genomic regions. A dashed curved line indicates the region from 3kb upstream of TSS to 3kb downstream of TTS; the numbers in parenthesis show the proportion occupied by each region in the genome. *Right*: Enrichment of normalized PADI2 reads in various genome regions relative to random distribution (* p-value <10^−2^; ** p-value <10^−3^; *** p-value <10^−4^). (**B**) Distribution of normalized PADI2 reads around the center of RNAP2 peaks in T47D cells. (**C**) RNAP2 and PADI2 occupancy across genes classified with increasing levels of expression. p-value was calculated by Wilcoxon-Mann-Whitney test in comparison to silent genes as indicated (** p-value < 10^−3^; *** p-value < 10^−5^).

To verify the specificity of the PADI2 occupancy, we performed PADI2 ChIP-qPCR in control (si*CTRL*) and PADI2 knockdown (si*PADI2*) over high (*SERPINA6*, c-*MYC*) and low expressed (*GSTT2*) genes and found that PADI2 depletion drastically decreased the levels (**Figure S6B**). PADI2 occupancy was significantly higher on genes downregulated by PADI2 depletion compared to those non-regulated (**Figure S6C**). Finally, RNAP2-ChIP followed by PADI2 re-ChIP revealed the association of RNAP2 and PADI2 at regulatory regions and gene bodies of the highly transcribed *SERPINA6*, c-*MYC* genes, but not in the low expressed *GSTT2* gene (**Figure S6D**). Thus, PADI2 seemed to be part of the transcription machinery in highly expressed genes.

### Citrullination of R1810 controls RNAP2 pausing

To investigate the role of R1810 citrullination in transcription, we analyzed RNAP2 occupancy by ChIP-qPCR in T47D cells prior and after PADI2 depletion. We observed accumulation of RNAP2 around the TSS of the highly expressed *SERPINA6, c-MYC, HMGN1* genes (**Figure 5A**). This effect is also dependent on R1810, as it is observed in T47D cells expressing only the α-amanitin resistant HA-tagged R1810A^r^ mutant form of RNAP2 in comparison with cells expressing only the WT^r^ RNAP2 (**Figure 5B**), suggesting absence of PADI2 mediated citrullination of R1810 leads to RNAP2 pausing. In T47D cells expressing only the R1810A mutant compared to cells expressing the WT^r^ RNAP2, ChIP-seq experiments showed remarkably high accumulation of RNAP2 at proximal promoters (**Figure 5C** *left panel*) and a corresponding change in the pausing index that correlated with the gene expression levels (**Figure 5C** *right panel*). Previously, Raji cells expressing R1810A also showed paused RNAP2 at proximal promoter for the 5% most highly expressed genes (Zhao et al., 2016).

**Figure 5 (With related Figure S7).**
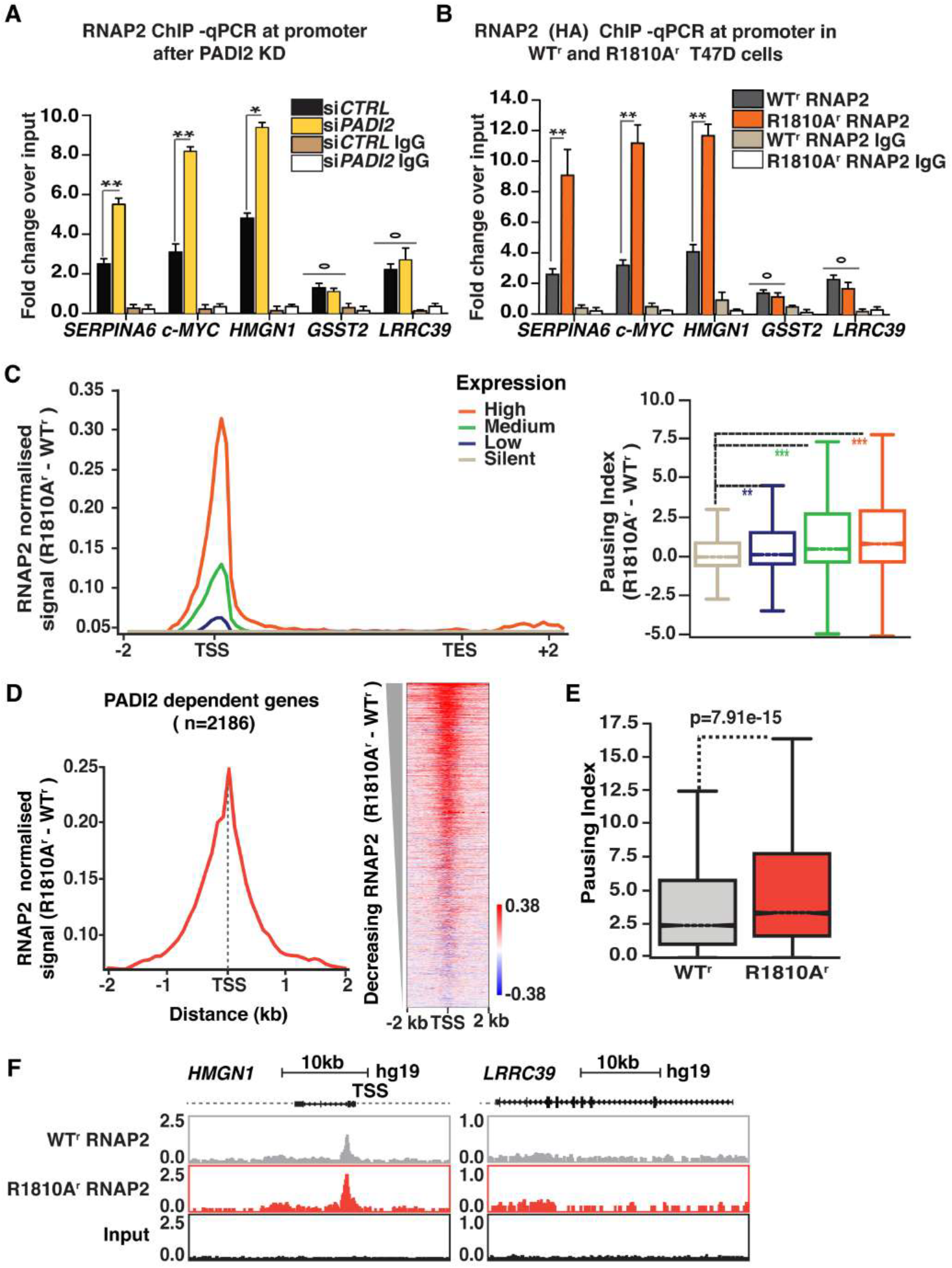
Cit1810 at CTD-RNAP2 regulates pausing in breast cancer cells. (**A-B**) RNAP2 ChIP qPCR assay performed in (**A**) T47D cells after PADI2 depletion with an antibody to RNAP2 (**B**) or in cells expressing only the HA-tagged wild-type (WT^r^) or R1810A^r^ mutant of RNAP2 with the HA antibody. Non-immune IgG was used as negative control. Y-axis: fold change over the input samples. (**C**) *Left*: Difference in RNAP2 density in T47D cells expressing HA-tagged R1810A^r^ mutant versus WT^r^ RNAP2 across genes classified by expression. *Right*: Pausing index of RNAP2 as indicated. (**D-E**) PADI2 dependent genes (n=2,186) showing (**D**) *Left*: average profile of difference in RNAP2 density (R1810A^r^ -WT^r^) around TSS. *Right*: heat map at TSS of genes ranked from highest to lowest RNAP2 density (R1810A^r^ -WT^r^). (**E**) Higher pausing index in cell expressing R1810A^r^ mutant as compared to WT^r^ form of RNAP2. (**F**) Browser snapshots showing RNAP2 occupancy for *HMGN1* & control gene *LRRC39* in cells expressing HA-tagged WT^r^ or R1810A^r^ form of RNAP2.

Focusing on PADI2-dependent genes (n=2186), we found a pronounced accumulation of RNAP2 around the TSS with a significantly increased pausing index in presence of R1810A mutant as compared to wild-type form of RNAP2 (**Figure 5D-F**). Similarly, genes downregulated in the presence of the R1810A^r^ RNAP2 mutant (n=939) also showed significantly higher pausing index as compared to cells expressing WT^r^ RNAP2 (**Figure S7B**). Thus, PADI2 depletion or absence of R1810 lead to RNAP2 accumulation on the promoters of highly expressed genes and to reduced level of S2P and S5P forms of RNAP2 (**Figure S7A**). We also found that genes upregulated upon depleting PADI2 or upon expressing the R1810A^r^ mutant RNAP2 exhibited lower pausing index (**Figure S7C-D**), suggesting that they do not need to overcome RNAP2 pausing to maintain their expression. In summary, we found that PADI2 citrullination of CTD R1810 is important for RNAP2 promoter pause release.

### Cit1810 at RNAP2-CTD recognized by P-TFEb

Citrullination is known to modulate functional protein-protein interactions (Tanikawa et al., 2018; Tanikawa et al., 2012; Vossenaar et al., 2003). We wondered whether the Cit1810 influences the interaction of RNAP2 with the components of P-TEFb complex CDK9 and CCNTI (Ccylin T1), which are required for RNAP2 pause release and productive elongation (Jonkers et al. 2015; Gressel et al., 2017). Immunoprecipitation of T47D cells extracts with a PADI2 antibody, precipitated CDK9 and CCNT1 (**Figure 6A**) and conversely a CDK9 antibody pulled down PADI2 (**Figure 6B**), indicating that PADI2 associates with the P-TEFb complex. Immunoprecipitation of extracts from PADI2 depleted cells (si*PADI2*) showed strong reduction in RNAP2 interaction with CDK9 and CCNT1 compared to control cells (si*CTRL*) (**Figure 6C**). In T47D or Raji cells expressing only the α-amanitin resistant HA-tagged WT^r^ RNAP2, CDK9 and CCNT1 were also immunoprecipitated with HA-tag antibody, and the interaction was significantly reduced in cells expressing the R1810A^r^ mutant RNAP2 (**Figure 6D-E**). Thus, PADI2 and R1810 are required for the association of RNAP2 with the CDK9-CCNT1 complex.

**Figure 6.**
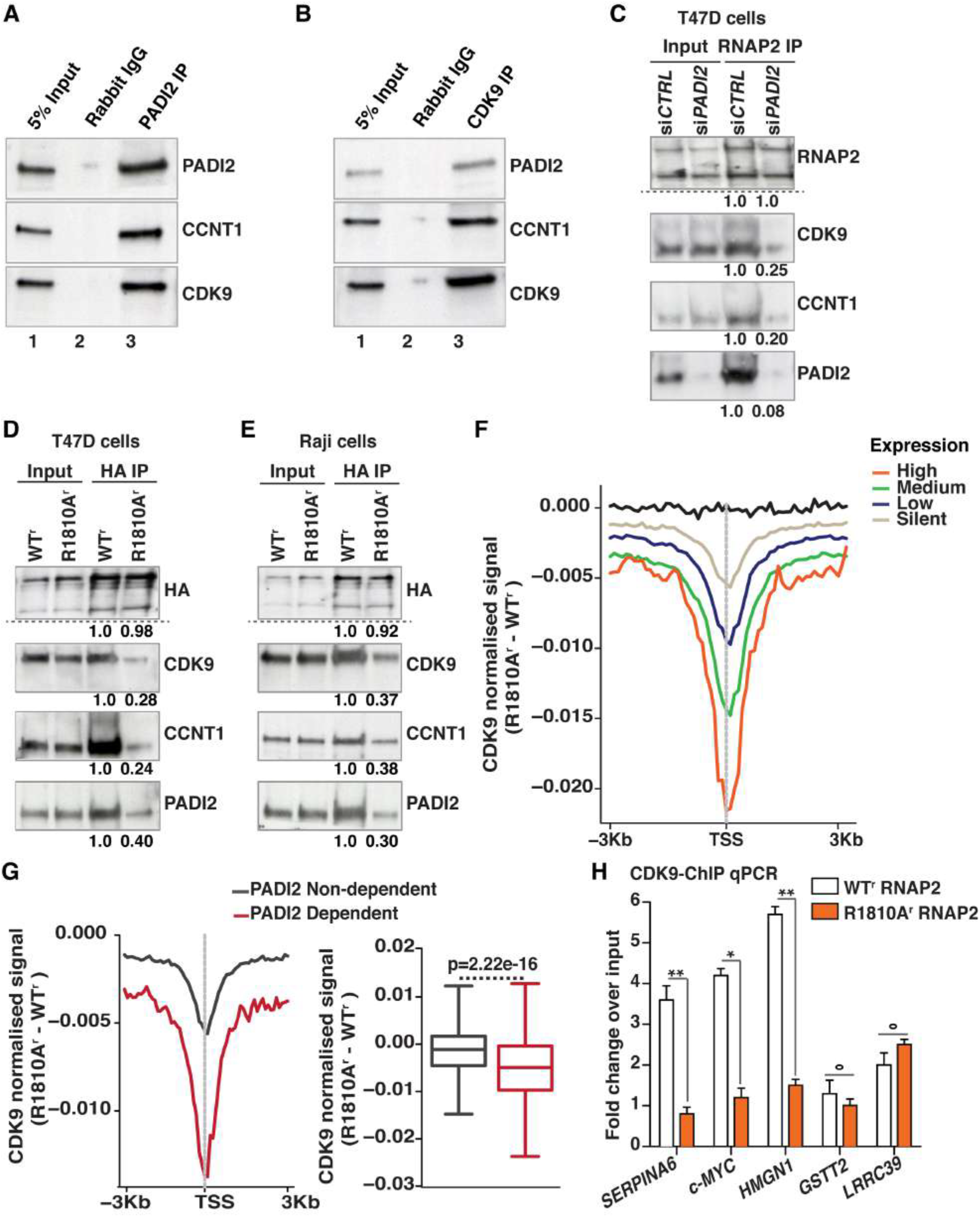
Cit1810 at RNAP2-CTD is recognized by P-TEFb. (**A-B**) Immunoprecipitation with (**A**) α-CDK9 (**B**) α-PADI2 or non-immune rabbit IgG of T47D extracts followed by western blot with the indicated antibodies. (**C**) Extracts from T47D cells in the presence (si*Ctrl*) or absence (si*PADI2*) of PADI2 were immunoprecipitated with α-total-RNAP2 followed by western blot for the indicated antibodies along with relative quantifications underneath. (**D-E**) Immunoprecipitation of extracts from (**D**) T47D (**E**) Raji cells expressing only the HA-tagged α-amanitin resistant WT^r^ or R1810A^r^ mutant of RNAP2 were precipitated with α-HA antibody and probed with the indicated antibodies. The relative quantification is shown underneath each gel. (**F**) Average profile of CDK9 density around TSS in ChIP-seq experiments using CDK9 antibody in T47D cells expressing the HA-tagged wild-type (WT^r^) or R1810A^r^ mutant of RNAP2 and difference (R1810A^r^ - WT^r^) across genes classified by expression level. Black line representing signal difference from random regions. (**G**) *Left panel*: Similar as in (F) for PADI2 dependent genes versus nondependent genes (siPADI*2*/siC*TRL*, FC<1.5; FC> 1/1.5, p-value > 0.05). *Right panel*: Box plot showing average difference in CDK9 profiles in PADI2 dependent versus non-dependent genes. p-value calculated by Wilcoxon-Mann-Whitney test. (**H**) Fold change over input in CDK9-ChIP qPCR assay on the promoter region of three PADI2 dependent genes (*SERPINA6, c-MYC, HMGN1*) and two non-dependent genes (*GSTT2, LRRC39*) in T47D cells expressing the HA-tagged wild-type (WT^r^) or R1810A^r^ mutant of RNAP2. Values are means ± SEM. * p-value < 0.05; ** p-value < 0.01; ° p-value > 0.05.

To explore whether R1810 is important for CDK9 recruitment to the promoter region of genes, we performed ChIP-seq with CDK9-antibody in T47D cells expressing only the α-amanitin resistant HA-tagged R1810A^r^ or the WT^r^ RNAP2. We found that CDK9 occupancy around the TSS decreased remarkably in the presence of R1810A mutant compared to wild-type form of RNAP2, particularly in highly expressed genes (**Figure 6F**), suggesting that the integrity of the R1810 is required for CDK9 recruitment. When we compared PADI2 dependent and non-dependent genes we also found a significant decrease of CDK9 occupancy in the presence of R1810A mutant as compared to wild-type form of RNAP2 (**Figure 6G**). We validated the ChIP-seq results by ChIP-qPCR and confirmed that mutation of R1810 significantly decrease the recruitment of the CDK9 to the promoter of PADI2 target genes *SERPINA6, c-MYC, HMGN1*, but not to the low expressed genes GSST2 and LRRC39 (**Figure 6H**). Altogether, this data supports the idea that PADI2 mediated citrullination of R1810 at RNAP2-CTD facilitates the recruitment of P-TEFb complex needed that promotes polymerase pause release and transcription of highly expressed genes involved in cellular proliferation.

### PADI2 unique residues contribute to citrullinate R1810 at RNAP2-CTD

We wondered about the reason why only PADI2 but not PADI3 or nor PADI4 carries out citrullination of R1810 at RNAP2-CTD. To address this issue, we used the published structure of the PADI2 (Slade et al., 2015) to model the attachment of RNAP2-CTD peptide encompassing R1810 (**Figure 7A**), and performed a similar structural modeling with the amino acids of PADI3. Our analysis revealed that the predicted binding score energies calculated with Rosetta were significantly lower for PADI2 in comparison to PADI3 (**Figure 7B**), indicating a higher affinity of PADI2 for the R1810 peptide.

**Figure 7.**
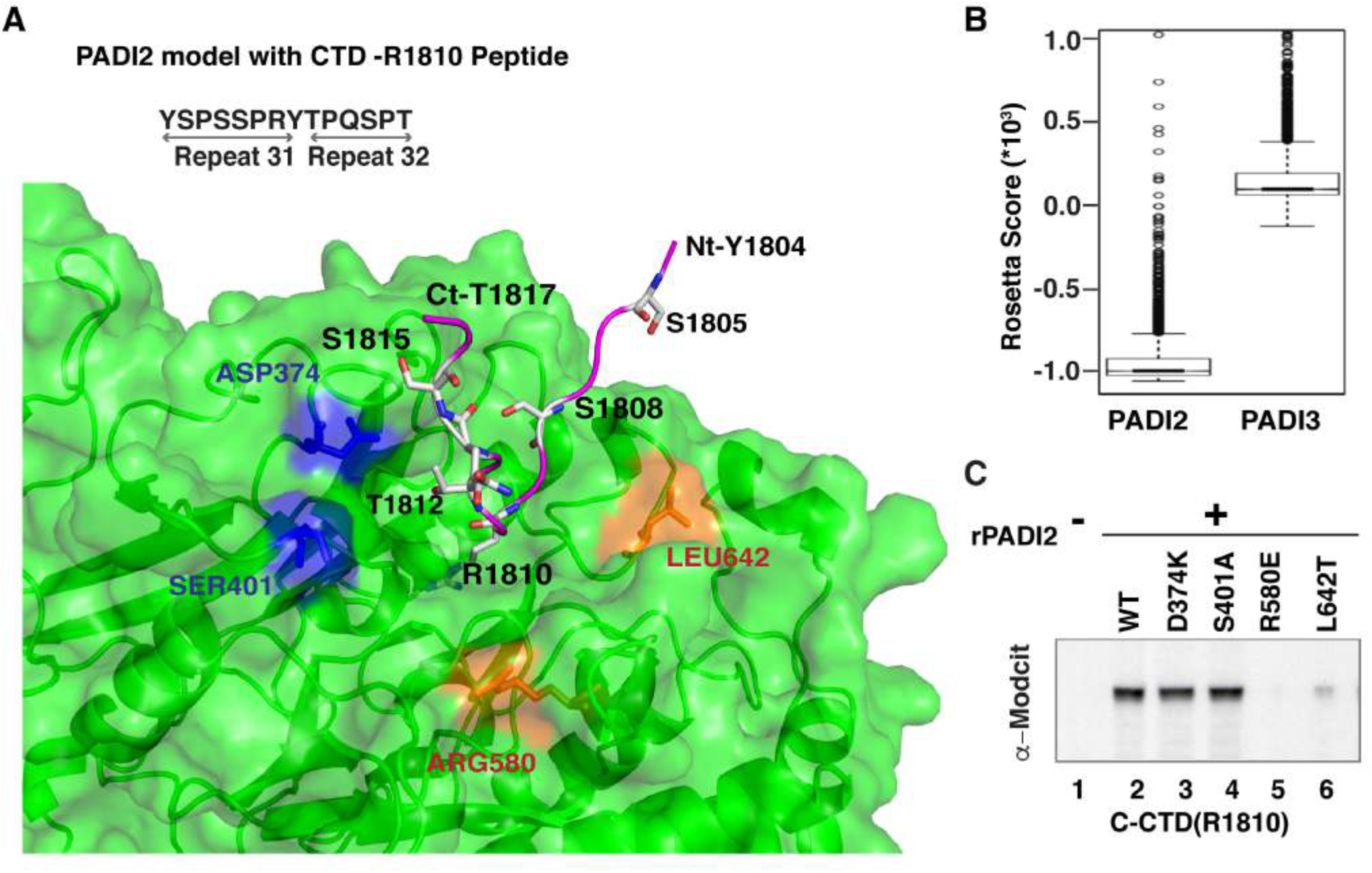
Illustration of structural model of PADI2 with R1810 at RNAP2-CTD. (**A**) Close-up of the peptide binding site of PADI2 shown in cartoon and surface representation (green) and R1810 CTD-RNAP2 peptide in ribbon (magenta) and stick representation of side-chains (colored by atom type: Oxygen: red; Nitrogen: blue; C: grey) of amino acid selected for mutation studies: non-conserved (ARG580, LEU642, shown in orange) and conserved (ASP374, SER401, shown in blue). (**B**) Box-plot showing the distribution of Rosetta Scores for the top 200 structural models performed for PADI2 and PADI3 proteins complexed with R1810 peptide. Central horizontal lines in the box mark the median and box edges of the first (Q1) and third (Q3) quartiles; top and bottom errors bars mark the Q1 and Q3 +1.5x interquartile range respectively; outliers are shown as empty circles. (**C**) *In vitro* citrullination immunoblot using the C-terminal CTD half containing R1810 as substrate, in absence (lane 2) or presence of recombinant PADI2 as wild-type (WT, lane 2) and PADI2 mutants of conserved residues (D374K and S401A lane 3 and 4) and of PADI2 unique residues (R580E and L642T, lane 5 and 6).

We next examined the PADI2 residues contributing most to the affinity for the R1810 peptide and analyzed their conservation in the PADI family. We found that nonconserved residues in PADI2 accumulate at the rim of the catalytic domain, most likely contributing to the R1810 peptide binding. We calculated Rosetta score for all PADI2 interface residues (including conserved and non-conserved) and ranked them according to conservation among PADI family members. Next, we choose PADI2 specific residues R580 and L642 and also two other PADIs conserved residues D374 and S401 that showed comparable low binding energy of PADI2 toward the R1810 peptide (**Figure 7A**), and introduced mutations that changed chemical properties of these four residues while maintaining the volume. Using GST C-CTD that encompasses R1810 as a substrate for *in vitro* citrullination assays, we found that the R580E and L642T mutations drastically reduced citrullination activity, whereas the D374K and S401A have a non-significant effect (**Figure 7C**), confirming the specificity of PADI2 unique residues for citrullination of R1810.

## DISCUSSION

In this study, we identify a novel post-translational modification of the RNAP2-CTD, namely the citrullination by PADI2 of R1810. This modification is coupled with the active form of RNAP2 and regulates the transcription of highly expressed genes involved in cell proliferation by mediating the interaction with P-TEFb and favoring RNAP2 pause release. We detected this modification initially in T47D breast cancer cells with a pan-citrulline antibody and confirmed it with Cit1810 antibody generated using a 13-mer CTD peptide centered on Cit1810. This modification is also present in several other breast cancer cell lines (MCF7, BT747, SKBR3), in HeLa cells, and also brain cancer cell lines (SK-N-SH and T98G), but is much less abundant in untransformed fibroblasts and breast epithelial cells. In breast cancer cells, PADI2 but not PADI3 or PADI4 specifically catalyze R1810 citrullination. Inhibition or depletion of PADI2 compromises transcription of thousands of highly expressed genes, as monitored by mRNA-seq analysis. Many of these PADI2-dependent genes are involved in key biological functions including RNAP2 transcription and cell proliferation. In chromatin RNA-seq, we found that PADI2 is mainly involved in active transcription, leading us to conclude that PADI2 participates in facilitating active transcription. However, in mRNA-seq analysis we also find genes up-regulated upon PADI2 depletion. Their up-regulation could be an indirect consequence of changes in the expression of PADI2-dependent genes or could be related to the need of R1810 dimethyation for proper transcription termination (Zhang et al., 2016). This remains to be directly demonstrated.

The previously reported asymmetrical (Sims et al., 2011) and symmetrical (Zhang et al., 2016) dimethylation of R1810 occurs mainly in hypo-phosphorylated RNAP2, as detected only after phosphatase treatment. In contrast, we find that cit1810 is preferentially associates with the transcriptionally engaged phosphorylated RNAP2. Depletion of the methyltransferases responsible for R1810 dimethylation, CARM1 and PRMT5, did not affect the expression of PADI2 dependent genes, and depletion of PADI2 did not change the expression of a broad range of snRNAs, the targets of CARM1. As PADI2 can not act on merthylate R1810, and citrullination will preclude methylation by arginine methyltrasferases, its seems that these are alternative types of modifications influencing different stages of transcription, as observed in other arginine residues that can undergo methylation and citrullination (Tanikawa et al. 2018; Cuthbert et al., 2004). This implies the dynamic nature of R1810 modifications, that change the docking surface for regulatory protein complexes to control various phases of transcription.

Our results of RNAP2 ChIP-seq show that PADI2 mediated cit1810 is required for RNAP2 pause release or high turnover (Krebs et al., 2017) at promoters of highly expressed genes that maintain cellular proliferation (Day et al., 2016; Gilchrist et al., 2010; Zeitlinger et al., 2007). In search for the molecular mechanism, we found that PADI2 interacts with the P-TEFb kinase complex, which is needed for RNAP2 pause release and productive transcript elongation. This interaction depends on R1810, supporting a function of cit1810 to facilitate transcription. Strikingly, by modeling how the structure of PADI2 could accommodate the R1810 CTD peptide, we identified PADI2-specific amino acid residues that are not conserved in other PADI family members, and when mutated interfere with citrullination of R1810 at RNAP2-CTD.

Although our results support a function of PADI2 mediated citrullination of CTD R1810 in transcription elogation, we cannot exclude an action of PADI2 on other substrates, including citrullination arginine 26 at histone H3 (H3R26), which could be involved in local chromatin decondensation and transcription activation (Zhang et al. 2012). Another intriguing open question concerns the implications of the proposed irreversibility of the citrullination reaction (Cuthbert et al., 2004; Wang et al., 2004). In the absence of enzymes that erase citrullination, alternative mechanisms may exit to replace the citrullinated RNAP2 before reaching transcription termination. Given that arginine-mediated interactions between intriniscally disordered protein domains, including the CTD of RNAP2, and RNAs or Poly(ADP-Ribose) are important in the formation of liquid droplets within the cell nucleus (Altmeyer et al. 2015; Hnisz et al. 2017; Harlen et al., 2017), we cannot exclude that citrullination of R1810 could also participate in modulating this interactions, and those influencing transcritional output. Further work will be required to investigate these possibilities.

Many elongation factors and kinases are implicated in the control of RNAP2 transcription pause release, a mechanism that controls the expression of genes involved in cancer progression and metastasis, like CDK9, MYC, JMJD6 (Zhang et al., 2017; Bywater et al., 2013; Miller et al., 2017). PADI2 is also found overexpressed in breast cancer (Cherrington et al., 2012) and other cancers (Guo et al., 2017). Indeed, we found that PADI2 depletion or mutation of R1810 reduced cell proliferation of breast cancer cells, by modulating cell cycle progression. Also, among *PADI* family members, only *PADI2* is overexpressed in breast cancer and other cancers and its overexpression correlates with poor prognosis. Thus, our finding opens the possibility that specific inhibition of citrullination at R1810-RNAP2 may represent a suitable drug target for combinatorial cancer therapy.

## AUTHOR CONTRIBUTIONS

P.S and M.B designed the study and strategy of the project; P.S performed the large majority of the experimental work; A.L performed immunofluorescence and image analysis. J.Q and J.C.C provided the bioinformatics analysis of all high throughput data. N.F.F and B.O performed structural analysis of data.R.H.G.W helped with Oncomine analysis. R.S and D.E generated stable Raji cells and helped with the R1810A^r^ mutant experiments. C.D.V and F.L.D performed ChIP sequencing, P.S and M.B discussed the results and wrote the paper. All other authors contributed to editing the manuscript.

## ACKNOWLEDGEMENTS

We thank David Bentley, for GST-CTD plasmids; Hiroshi Kimura, for RNAP2 S2P/5P monoclonal antibodies. We thank CRG Genomics, Protein Technology and the Advanced Light Microscopy facilities for all technical support. We are grateful to all the members of the chromatin and gene expression lab for useful suggestions. We acknowledge Juán Valcárcel, Guillermo P. Vicent, Enrique Vidal Ocabo, and Gwendal Dujardin from CRG for constructive criticism and advice during the course of this work. P.S. was supported by a Novartis fellowship and Beatriu de Pinós fellowship co-founded by Marie Curie Action (2013 BP_B 00061). Our work supported by Spanish MEC (SAF2013-42497), the Catalan Government (AGAUR 2014SGR780) and the European Research Council Synergy Grant “4DGenome” (609989). We acknowledge the support of the Spanish Ministry of Economy and Competitiveness, ‘Centro de Excelencia Severo Ochoa” and the CERCA Programme / Generalitat de Catalunya”.

## Supplementary Figures

**Figure S1 (Related to Figure 1).**
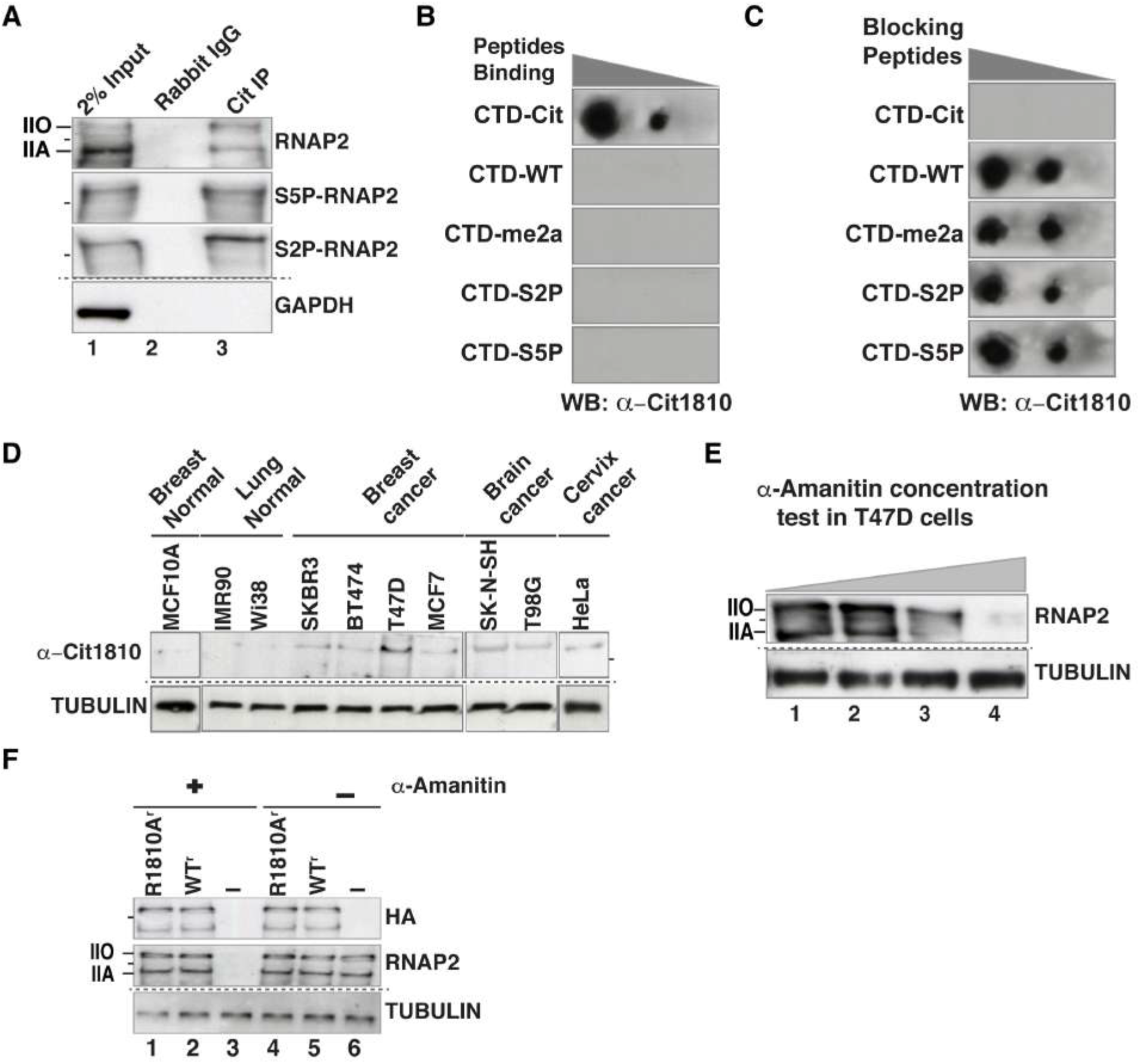
Specificity of the anti-Cit1810 RNAP2 antibody. (**A**) T47D cells extracts immunoprecipitated with pan-citrulline antibody and analyzed by western blot using antibodies against total-RNAP2, S5 and S2 phosphorylated forms of RNAP2. The dotted line indicates a separate gel (see methods) and lines on the left mark the migration of the 250 kDa size marker. (**B**) Dot blot showing the specificity of α–Cit1810, performed with 1, 0.2 and 0.04 μg of mentioned RNAP2-CTD peptides encompassing R1810 with specific post-translational modifications. Only the peptide bearing the Cit1810 modification were specifically recognized by α-Cit1810. (**C**) The specificity of α-Cit1810 was also confirmed by the peptide competition assay. α-Cit1810 was incubated with 2μg of each of the indicated peptides for 30 minutes at 4°C prior to probing the membrane with the Cit1810 peptide. (**D**) Cell extracts from indicated cell lines were probed in western blot with with α-Cit1810 and tubulin antibody (**E**) Titration of the concentration of α–amanitin needed to deplete endogenous RNAP2 in T47D cells. Cells were incubated with increasing concentrations of α–amanitin (0, 2, 4 and 6μg/ml; *lanes 1 to 4*) for 12h before preparing nuclear extracts. The extracts were probed in western blot with an antibody against total RNAP2; Tubulin was used as loading control. (**F**) Western blot of extracts from cells depleted of endogenous RNAP2 by preincubation with 6μg/ml α–amanitin, and transfected with empty vector (−) or with expression vectors for WT^r^ or the R1810A^r^ mutant HA-tagged recombinant RNAP2 carrying an additional mutation that makes the enzyme resistant to α–amanitin (see Method). The western blots were probed with antibodies against the HA tag or total-RNAP2; Tubulin was used as loading control.

**Figure S2 (Related to Figure 2).**
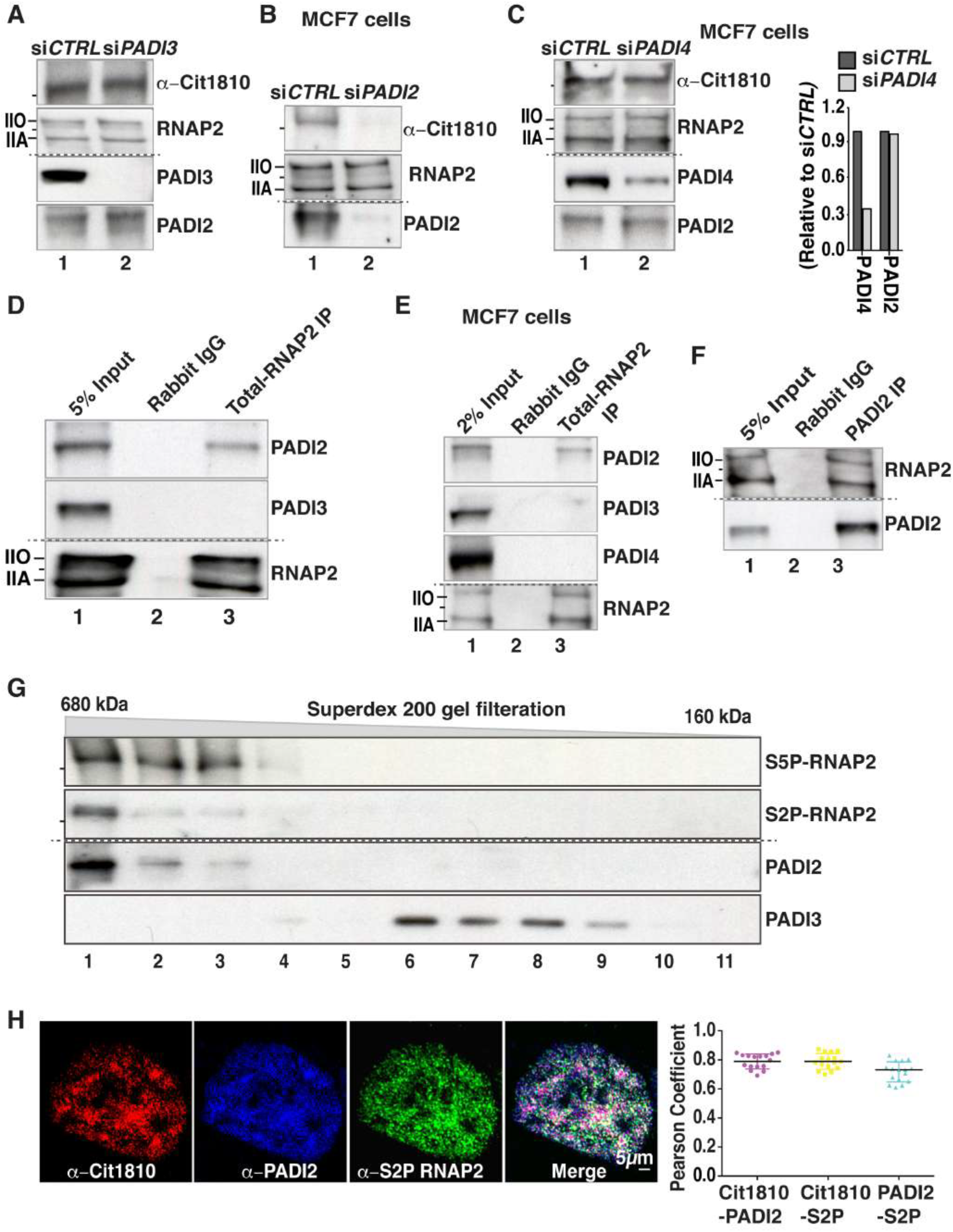
PADI2 interacts with CTD-R1810 in *vivo* and *in vitro*. (**A**) Western blot of extracts from T47D cells transfected with siRNA control (si*CTRL)* or siRNA against *PADI3* (si*PADI3)* and probed with the indicated antibodies. (**B**) Western blot of extracts from MCF7 cells transfected with si*CTRL* or si*PADI2* and probed with the indicated antibodies. (**C**) Western blot of extracts from MCF7 cells transfected with siRNA *CTRL* or si*PADI4* and probed with the indicated antibodies. The bar plot represents the total intensity of PADI4 and PADI2 band quantitated by Image J after PADI4 depletion. (**D-E**) Immunoprecipitation of (**D**) T47D (**E**) MCF7 extracts with an α-total-RNAP2 or with non-immune rabbit IgG followed by western blot with the indicated antibodies. (**F**) Immunoprecipitation of T47D extracts with α-PADI2 or non-immune rabbit IgG probed with indicated antibodies. (**G**) Fractionation of T47D cells extract using a Superdex 200 gel filtration column and analysis of the eluted fractions by western blotted with the indicated antibodies. (**H**) *Left*: Representative super-resolution immunofluorescence images of T47D cells using triple labeling with α-Cit1810 (red) in combination with α-S2P-RNAP2 (green) and α-PADI2 (blue). Plot representing the mean Pearson correlation coefficient of several individual cells (n=16), for PADI2 with Cit1810 (purple), for S2P-RNAP2 with Cit1810 (yellow) and for PADI2 with S2P-RNAP2 (cyan). *Right*: Person correlation coefficient presented a value of mean ± SEM. Note Cit1810-RNAP2 showed co-localization with PADI2 as well as with S2P-RNAP2 (shown as white in the merged image).

**Figure S3 (Related to Figure 2).**
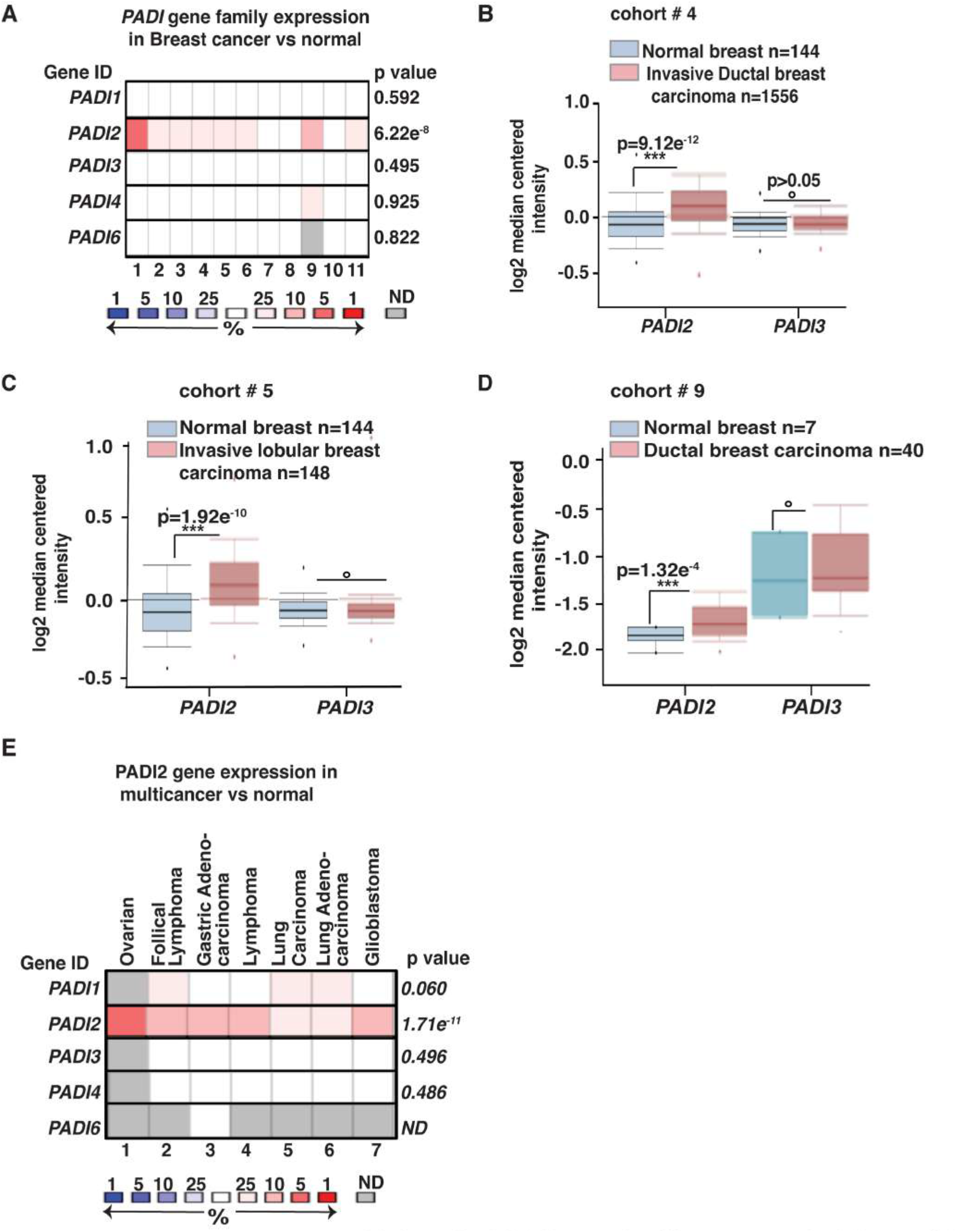
*PADI2* overexpressed in breast and other cancers. (**A**) *PADI* gene family expression in breast cancer cohorts (n=2155 patients). Each column represents a breast cancer cohort. Scale on the bottom is the best gene rank percentile; color indicated overexpression (red) and underexpression (blue) compared to normal tissue from same study. The p-value for a gene is its p-value for the median-ranked analysis. Note that only over expression of *PADI2* but no other *PADI* family member is significant. (**B-D**) *PADI2* and *PADI3* expression level were analyzed in various subtypes of breast cancer compared to normal breast: (**B**) Cohort-4, normal breast versus invasive ductal breast carcinoma; (**C**) Cohort-5, normal breast versus invasive lobular breast carcinoma; (**D**) Cohort-9, normal versus ductal breast carcinoma. Y-axis represents log2 median intensity. The p-value indicates the significance of difference, ° for p > 0.05. (**E**) *PADI* gene family expression in 7 different cancer cohorts, represented by each column. The p-value for a gene is its p-value for the median-ranked analysis. Scale at the bottom is as (**Figure S3A**).

**Figure S4 (Related to Figure 2).**
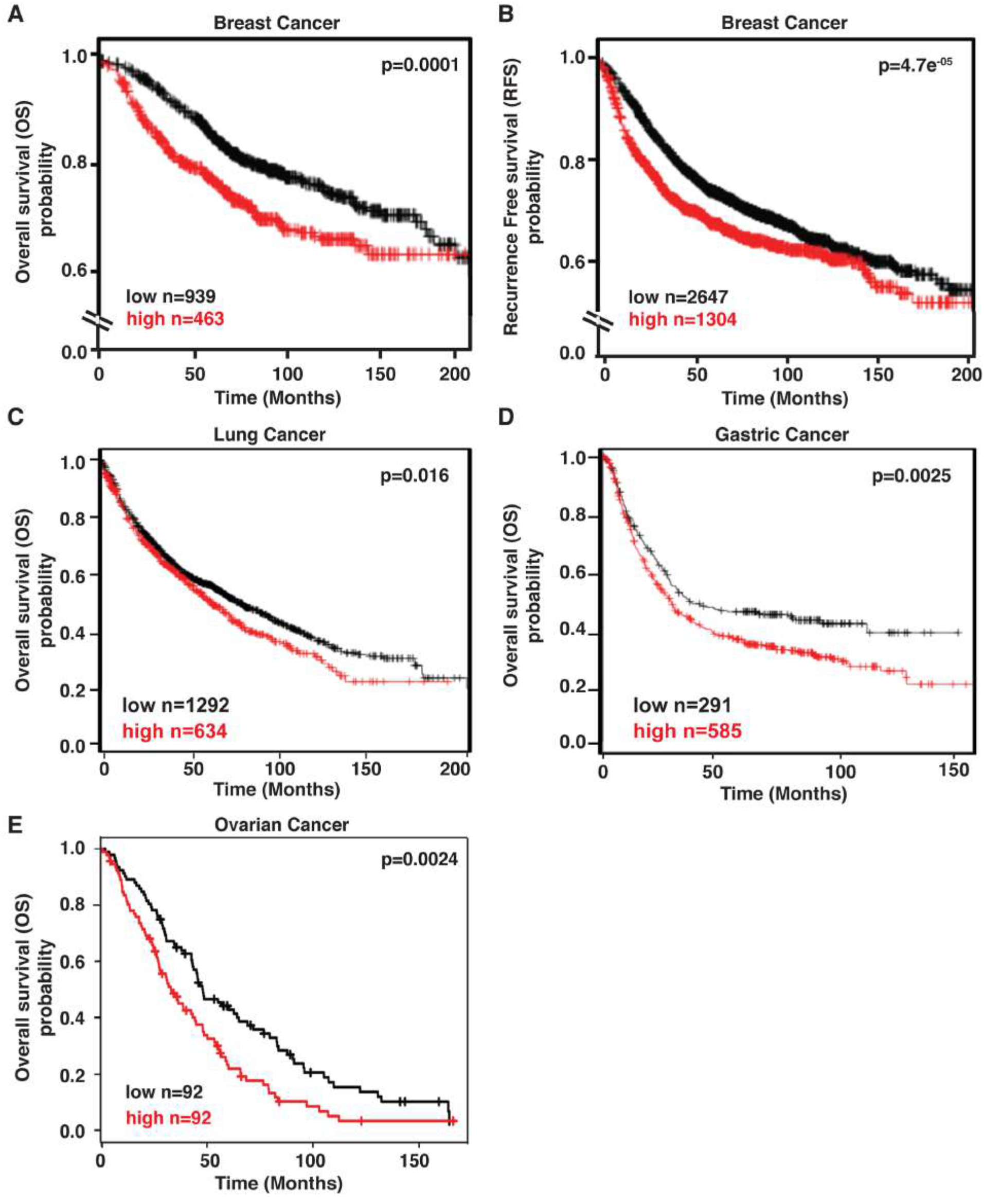
Elevated levels of *PADI2* in patient samples associates with poor prognosis. (**A-B**) Kaplan-Meier survival graph segregated according to *PADI2* expression, high (> 75% percentile) or low (<25% percentile): (**A**) overall survival; (**B**) recurrent free survival for breast cancer patients. (**C-E**) Kaplan-Meier survival graph showing overall survival for (**C**) lung cancer (**D**) gastric cancer and (**E**) ovarian cancer (GSE26712). The number of patients and the p-value of the comparison are indicated.

**Figure S5 (Related to Figure 3).**
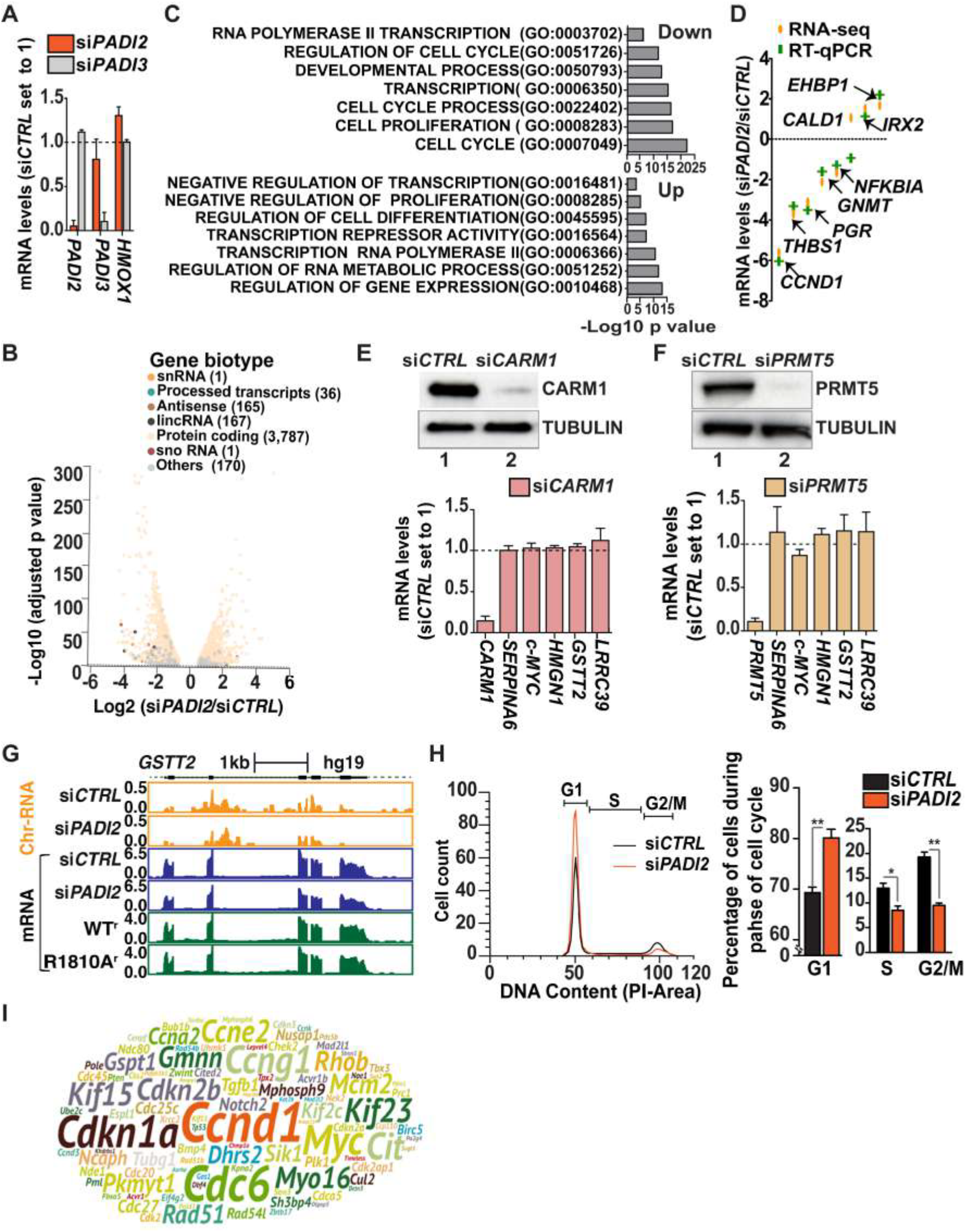
PADI2 depletion affects genes involved in cell cycle progression. (**A**) Changes in *PADI2, PADI3* and *HMOX1* mRNA quantified by RT-qPCR in T47D cells transfected with si*PADI2* (orange bars) or si*PADI3* (grey bars). mRNA levels were normalized to a *GAPDH* mRNA expression level, which was not affected in these experimental conditions, and are presented as a ratio to levels in cells transfected si*CTRL*. Values are the mean ± SEM of at least three biological replicates as in other plots in the figure. (**B**) Volcano plot showing indicated gene biotypes of differentially expressed genes determined from polyA-RNA sequencing as shown in Figure 3A. The X-axis represents log2 expression fold changes (FC) and the Y-axis represents the adjusted p-values (as −log10). (**C**) Gene set enrichment analysis (GSEA) for biological processes and molecular functions. Seven representative processes are presented. The X-axis shows the −log10 transformed p-values. GO, Gene Ontology. (**D**) The mRNA-seq data validated by RT-qPCR for indicated genes. Y-axis shows the fold change over si*CTRL*, orange color indicates data from duplicate RNA-seq and green color from RT-qPCR (mean ± SEM from at least three experiments). (**E**) Knockdown of *CARM1* mRNA (>80% depletion relative to si*CTRL*) did not significantly affect PADI-dependent gene transcripts. Data are shown as fold change over si*CTRL* and represent mean ± SEM. Extracts from si*CTRL*and si*CARM1* transfected cells were probed with indicated antibodies to estimate CARM1 depletion (>80%). (**F**) Knockdown of *PRMT5* mRNA (>80% relative to si*CTRL* mRNA) did not significantly affect PADI-dependent gene transcripts. Extracts from si*CTRL* and si*PRMT5* transfected cells were probed with indicated antibodies to estimate PRMT5 depletion (>90%). (**G**) Browser snapshots of control *GSTT2 (*related to **Figure 3D**) gene in T47D cells showing chromatin associated RNAs sequencing profile (orange) and mRNA sequencing profiles after *PADI2* knockdown (blue) and expressing WT^r^ and R1810^r^ mutant form of RNAP2 (Green). Scale is indicated on the top of the gene. (**H**) *Left*: Cell cycle profile of T47D cells transfected with si*CTRL* (black) and si*PADI2* (orange) obtained by propidium iodide labeling followed by fluorescence-activated cell sorting (FACS) analysis. *Right*: Histogram showing the percentage of cells during cell cycle phases. Data presented as mean ± SEM from three biological replicates. p-value was calculated by student’s t test, * p-value < 0.05, ** p-value < 0.001. (**I**) Word cloud of all the PADI2 dependent genes represented in Figure 3H; Size of genes indicates an extent of down-regulation after PADI2 depletion as measured by the log2 fold change si*PADI2*/si*CTRL*.

**Figure S6 (Related to Figure 4).**
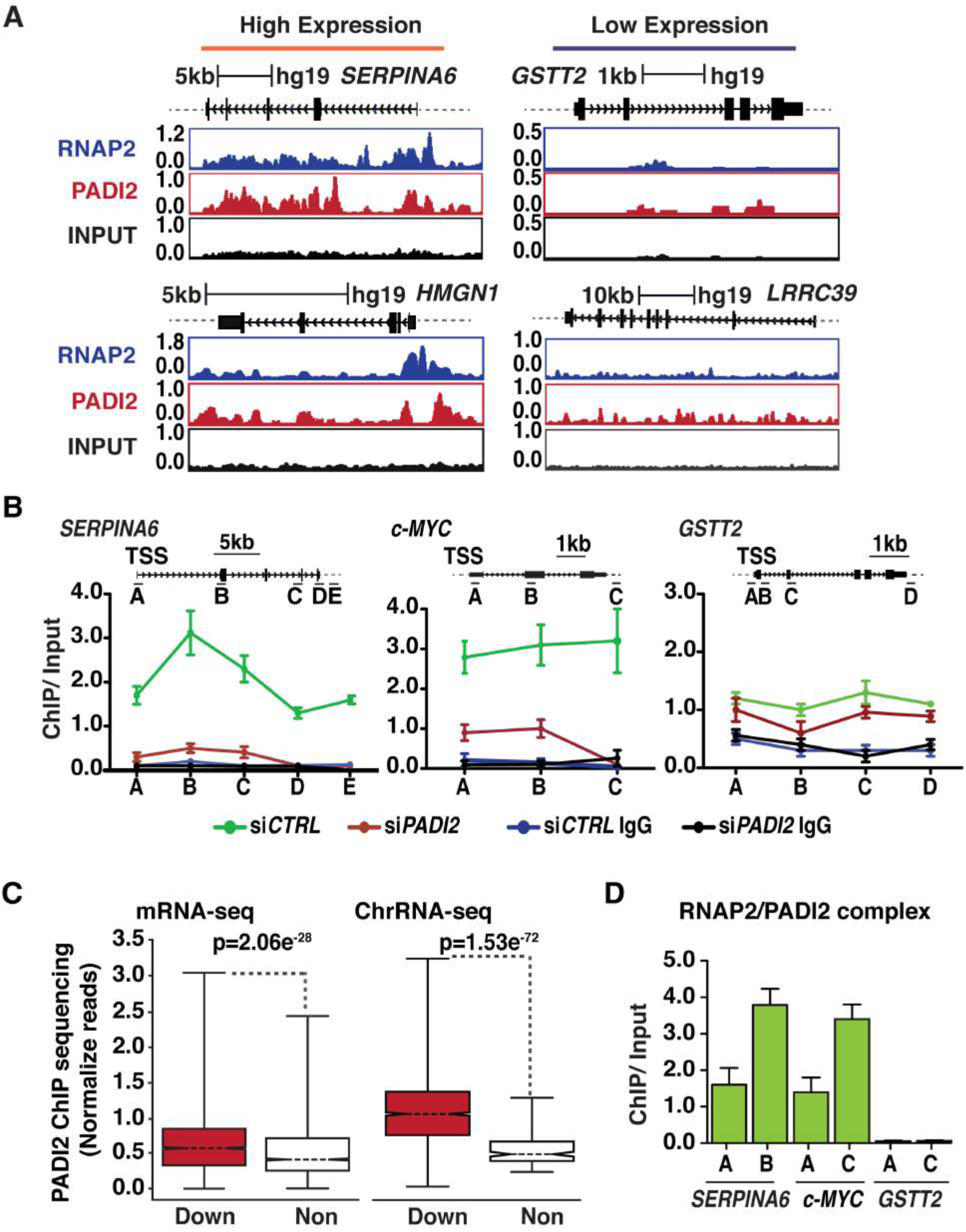
RNAP2 and PADI2 co-localize on highly transcribed genes. (**A**) Browser snapshots representing the PADI2 and RNAP2 occupancy on highly transcribed genes *SERPINA6* and *HMGN1* versus lowly expressed genes (*GSTT2* and *LRRC39*. A scale is shown on the top for each gene. Y-axis: reads per million (RPM). (**B**) PADI2 occupancy monitored along the gene bodies by PADI2 ChIP assay in T47D cells transfected with si*PADI2* or si*CTRL* followed by qPCR along the gene bodies of two highly transcribed genes (*SERPINA6, c-MYC*) and a low transcribed gene (*GSTT2*). Non-immune IgG was used as negative control. Y-axis: PADI2 enrichment over input samples. Data represented as mean ± SEM from at least three biological replicates as in other plots of the figure. Top: basic gene structure and scale with the positions of the amplicons. (**C**) PADI2 ChIP-seq normalized reads in a window from 3kb upstream of TSS to 3kb downstream of TTS on genes downregulated and nonregulated genes after PADI2 depletion in mRNA-sequencing (*left*) and ChrRNA-sequencing (*right*). Downregulated genes showed significantly higher PADI2 recruitment both from mRNA-seq and ChrRNA-seq as indicated, p-value was calculated by Wilcoxon-Mann-Whitney test. Box represents the interquartile range; Whisker extends the box to the highest and lowest values. A line across each box indicate the median value. (**D**) RNAP2 ChIP followed by PADI2 re-ChIP assay performed in T47D cells to examine co-localization on promoter regions (A primer) and exons (B or C primers) of two highly transcribed genes (*SERPINA6, c-MYC*) and a low transcribed gene (*GSTT2*).

**Figure S7 (Related to Figure 5).**
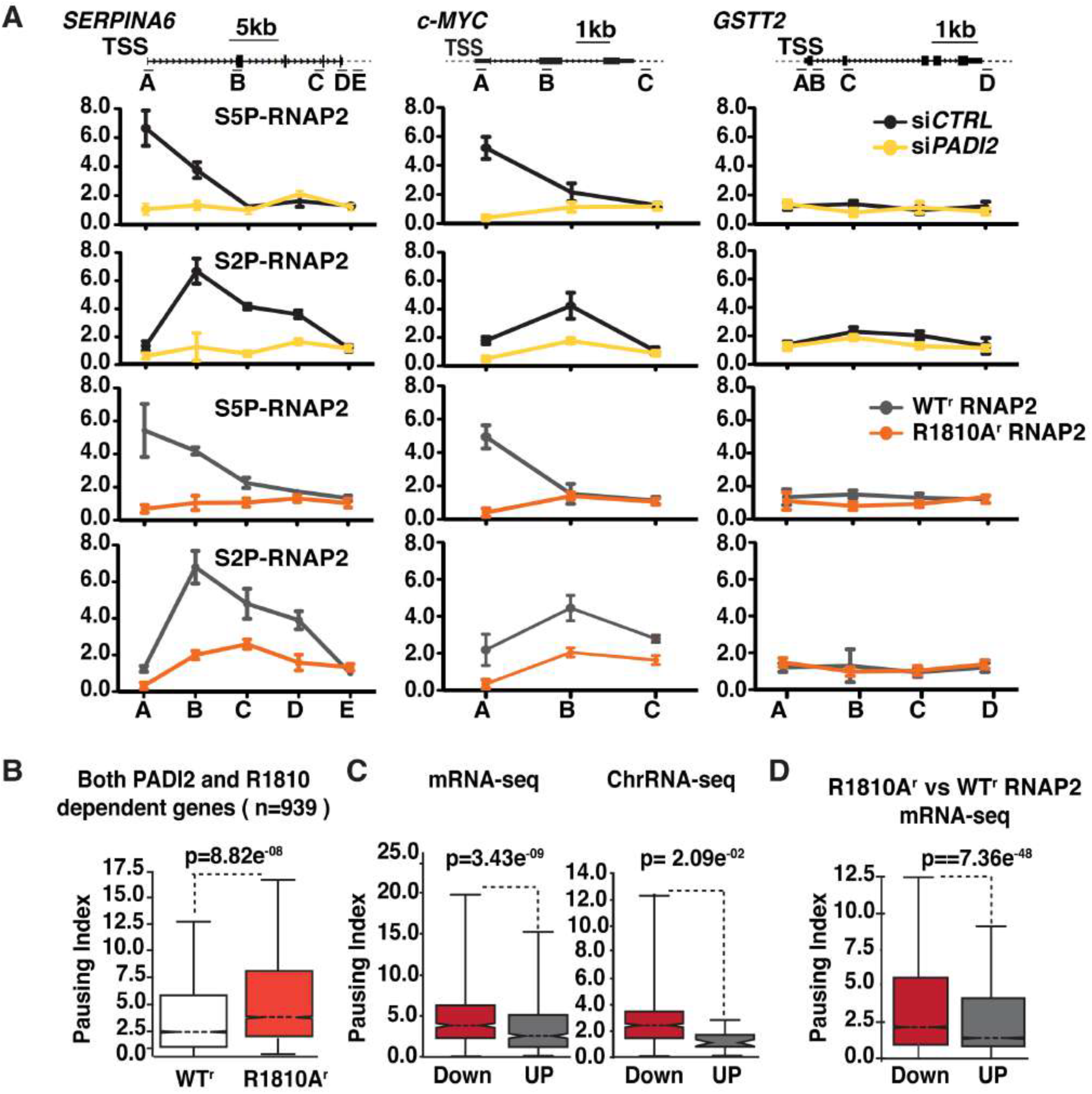
PADI2 directed citrullination of R1810 regulates pausing of highly transcribed genes. (**A**) S5P and S2P RNAP2 occupancy was monitored along the gene bodies by performing ChIP assay in T47D cells transfected with si*CTRL or* si*PADI2 (black or yellow)*, or T47D cells expressing only WT^r^ or the R1810A^r^ mutant of RNAP2 (grey or orange) followed by quantitative PCRs along the gene bodies. Y-axis: relative enrichment normalized to total RNAP2; values are means ± SEM. *Top*: For each gene, the basic gene structure and the position of the amplicons are indicated. (**B**) Higher pausing index in the cell expressing R1810A^r^ mutant as compared to WT^r^ form of RNAP2 calculated for shared 939 genes downregulated after PADI2 depletion (2,186) and R1810 dependent (1392 genes found significantly downregulated by considering FC < 1/1.5 and p-value < 0.01, after expressing R1810A^r^ mutant as compared to WT^r^ RNAP2) in T47D. (**C-D**) Box plot of the pausing index of (**C**) top 25% down-regulated and up-regulated genes after PADI2 depletion from mRNA-seq (*left*) and ChrRNA-seq (*right*). (**D**) Downregulated (n=1392) and upregulated (n=1372) genes obtained from replicate mRNA-seq performed in T47D cells expressing R1810A^r^ mutant as compared to WT^r^ form of RNAP2. p-value was calculated by Wilcoxon-Mann-Whitney test. Each box represents the interquartile range; Whisker extends the box to the highest and lowest values. A line across each box indicate the median value.

